# Multimodal fertility cues in chimpanzees: How body odours complement sexual swellings

**DOI:** 10.64898/2026.05.21.726750

**Authors:** Marlen Kücklich, Madita Zetzsche, Sofya Dolotovskaya, Jakob W. Siepmann, Lydia Schmidt, Claudia Wiesner, Brigitte M. Weiß, Anja Widdig

**Author notes:** These authors contributed equally to this work and share second authorship (Madita Zetzsche, Sofya Dolotovskaya).

## Abstract

To attract mating partners, female mammals communicate their reproductive status through one or multiple sensory modalities, providing redundant or complementary information. Chimpanzees (*Pan troglodytes*) are an excellent model for studying multimodal communication. Exaggerated sexual swellings of females serve as a visual proxy for ovulation but increased male mating interest during maximum swelling suggests that olfactory cues may pinpoint fertility more accurately than the swelling alone. Here, we combined gas chromatography–mass spectrometry, hormonal analyses, and bioassays to examine (1) whether chemical composition of female anogenital odours changes during the fertile period, and (2) whether males are able to detect these changes. Our results suggest that, in addition to prominent olfactory changes associated with swelling stages, chemical cues provide complementary information regarding the timing of the fertile window. These changes, however, are minor compared to those related to swelling stages. Male behavioural responsiveness in bioassays was too low to draw conclusions regarding their ability to detect these subtle shifts when presented with a chemical cue only. Overall, our findings support the existence of a multimodal fertility cue in chimpanzees, wherein visual signals are complemented by subtle olfactory changes indicating the timing of the fertile period.

## INTRODUCTION

Mating success is a fundamental part of individual fitness. To attract mating partners, female mammals communicate their reproductive status through various sensory modalities, including vision, acoustics, and/or olfaction (1,2). Sexual selection exerts strong pressures on traits that improve inter-sexual communication (2). Sensory modalities may act alone, but highly social, group-living animals often engage in multimodal communication (3,4). For example, many species of birds show complex visual displays during singing (5–7), and some bats combine vocalizations with fanning odours toward potential mates (8).

Primates are remarkable in the diversity of sensory modalities used to communicate the reproductive status (9). Female primates can use visual cues, such as sexual swellings in chimpanzees (*Pan troglodytes*) (10) and olive baboons (*Papio anubis*) (11) or sexual skin colour in rhesus macaques (*Macaca mulatta*) (12,13). Some primates use auditory cues, such as copulation calls in Barbary macaques (*Macaca sylvanus*) (14), or behavioural cues, such as solicitations and other proceptive behaviours in black howler monkeys (*Alouatta pigra*) (15) and long-tailed macaques (*Macaca fascicularis*) (16). Finally, female primates use olfactory cues, such as scent marks in common marmosets (*Callithrix jacchus*) (17) and ring-tailed lemurs (*Lemur catta*) (18). These cues can be exhibited in various combinations, whereby different components may transmit redundant or complementary information (19). Some indicators of reproductive status may constitute *signals*, i.e., traits that evolved to communicate fertility-related information to conspecifics, or *cues*, i.e., traits that did not evolve for communicative purposes, e.g., being by-products of hormonal changes (20).

Whereas visual and acoustic cues of fertility have been studied well in non-human primates, recent studies also highlight the importance of smell, a modality which has long been neglected in primate communication studies (21–23). Chemical analyses of female body odours demonstrated differences between cycling and non-cycling females (ring-tailed lemurs (18)), different phases of the menstrual cycle (common marmosets (17); Japanese macaques, *Macaca fuscata* (24); olive baboons (25)), and stages of sexual swelling (chimpanzees (26)). Some studies identified specific chemical substances that are related to fertility (17,24–26). Behavioural studies further demonstrated that female body odours can convey fertility-related information to males. In several observation studies, olfactory inspection of the female anogenital region by males increased during the fertile period (olive baboons and chacma baboons, *Papio ursinus* (27,28). In behavioural tests, males of several primate species reacted to body odours collected from fertile females with increased exploratory and arousal behaviours (common marmosets (17,29); cotton-top tamarins, *Saguinus oedipus* (30)), as well as with an increase in testosterone levels (common marmosets (31); stumptailed macaques, *Macaca arctoides* (32)).

Chimpanzees (*Pan troglodytes*) represent an excellent model for investigating whether olfaction integrates with other sensory modalities to communicate fertility multimodally. Female chimpanzees exhibit several cues that may indicate fertility with different levels of precision. Females have exaggerated sexual (anogenital) swellings, which vary in size across the menstrual cycle phases but are not precisely aligned with them (33). The period of maximal swelling lasts for about one third of the menstrual cycle (10-12 days on average, with the length of the menstrual cycle averaging 36 days) and includes parts of the follicular and periovulatory phases (33–35). Ovulation occurs within the period of maximal swelling, and although the size of the swelling slightly increases as ovulation approaches, the precise ovulation day cannot be determined from the swelling size alone (36,37). Females also exhibit proceptive and resistance behaviours at different rates during fertile and non-fertile periods, with higher proceptivity and lower resistance to preferred males during the fertile period compared to non-fertile period (38). Finally, females produce copulation calls, although their rates or acoustic structure do not vary between fertile and non-fertile periods (39,40). None of these cues, however, provides precise information about the fertile period, and it has been suggested that concealing ovulation allows females to prevent monopolization by dominant males and thereby confuse paternity and reduce infanticide risk (40,41). In line with this, although copulations occur most frequently during maximal swelling, mating also occurs outside this period (42).

Despite this presumed imprecision of fertility cues, observations suggest that male chimpanzees get more precise information about fertile days from olfactory cues than from visual cues (swelling size) alone. First, males sniff at females more often during maximal swelling (43). Second, males display increased mating interest during the periovulatory period compared to other days of maximal swelling (36,44). As male competition over the females is intense, with high-ranking males gaining most of the copulations (45–47), males should benefit from using precise fertility indicators, as it would allow them to maximize mating efforts on fertile females. A potential mechanism for olfactory fertility cues may involve cycle-dependent hormonal changes. Since levels of steroid hormones vary in relation to sexual swelling stage as well as menstrual cycle phase (48,49), hormone-related olfactory changes may correspond to both processes.

Previous studies in chimpanzees have found that some volatile fatty acids in female vaginal secretions change in levels across the menstrual cycle (50,51). A more recent study, using gas chromatography–mass spectrometry (hereafter, GC–MS) to analyse the chemical composition of female body odours (from torso and limbs) across stages of sexual swelling, showed that chemical composition of odours differs between swelling stages (26). However, it remains unclear whether chemical profiles of body odours vary between menstrual cycle phases, in addition to variation across swelling stages, and whether this variation provides more precise information about fertility than the swelling size alone. Here, we used information not only on swelling stages, but also on the menstrual cycle phases, obtained from hormonal analyses, to investigate both of their effects on the chemical composition of female anogenital odours. We also investigated male perception of female anogenital odours by presenting males the odours from unfamiliar females in various menstrual cycle phases. Our study had two aims: (1) to unravel the impact of sexual swelling stage and menstrual cycle phase on the chemical composition of anogenital odours of female chimpanzees; and (2) to examine whether male chimpanzees can distinguish between the anogenital odours from females in different phases of the menstrual cycle. We predicted that the chemical composition of female odours will vary across the menstrual cycle phases, in addition to the variation between the swelling stages, and therefore provide more precise information on fertility than the swelling alone, and that males will be able to detect these differences.

## METHODS

### Sampling

Samples were collected by L.S. in June and July 2022 at Zoo Dudley (Dudley, UK) from five naturally cycling female chimpanzees aged 27 to 47 years (mean ± SD: 38.8 ± 9.1) living in a social group of seven females. The animals were trained using positive reinforcement to voluntarily participate in sampling. Samples were collected over one complete menstrual cycle (mean ± SD: 34 ± 2.5 days), from the onset of increasing swelling to the next. We differentiated four swelling stages: flat, increasing, maximum, and decreasing, following (34) (Fig. 1).

**Fig. 1.**
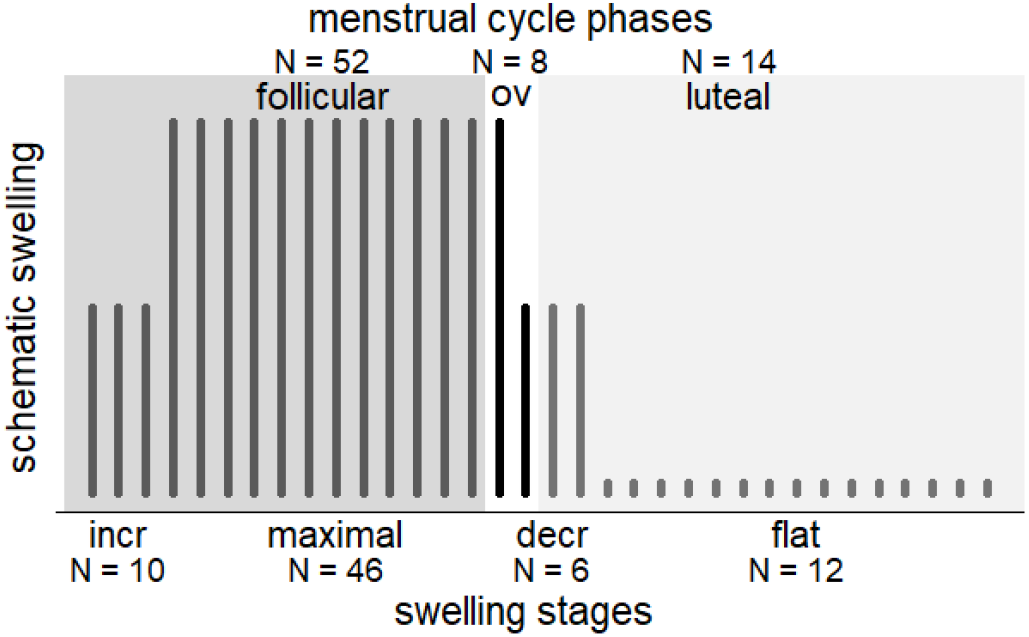
Schematic representation of swelling stages using dark bars (small = flat, medium = increasing/decreasing, large = maximum swelling) reflecting the average cycle length over all females (increasing: 3 days, maximal: 13 days, decreasing: 3 days, flat: 15 days). Sample distribution across the menstrual cycle phases is shown above the graph and across the swelling stages below. The periovulatory window (“OV”) is shown as white area.

We collected *saliva samples for hormone analysis* using synthetic swabs (Salivette® Cortisol, Sarstedt, Germany). Females were trained to take swabs directly with their mouth, chew for at least 10 sec, and return the chewed swab by presenting it between the teeth. The saliva was retrieved by manually centrifuging the swabs. Swabs were stored at −18 °C during sample collection and at −80 °C for storage. In parallel, we collected *odour samples for chemical analysis and bioassays* from the anogenital region since a previous study found that males mainly sniffed at this area (43). Animals were trained to present their anogenital region to the grid of the enclosure for sampling. We used different methods for GC-MS analysis (optimized for volatile preservation) and bioassays (optimized for volatile release during bioassays), both successfully used in this combination in previous studies (17,52).

*Odour samples for chemical analysis* were collected using preconditioned thermal desorption (TD) tubes (stainless steel, 1/4 in. × 3 1/2 in., Supelco, Bellefont, USA) containing Tenax TA (0.095 g) and XAD-2 (0.21 g, Sigma-Aldrich, Steinheim, Germany, see (17) for details) from close to the skin (about 1 – 2 cm away). A BiVOC2 pump (Umweltanalytik Holbach GmbH, Germany) was used to draw 0.5 L air through pre-cleaned TD tubes with a flow of 1 L/min for 30 sec. Afterwards, TD tubes were closed with Swagelok Brass caps (1/4 in., Supelco, Bellefont, USA) and stored at room temperature. We then excluded samples that fell between swelling stages or cycle phases (i.e., first day of maximal swelling, three days before and one day after periovulatory window), resulting in a total of 74 samples (see Fig. 1 for sample distribution across cycle phases and swelling stages; 12-17 samples per female, mean ± SD: 14.8 ± 2.2). We included three types of blank samples: environmental blank samples, collected in front of the enclosures in the absence of animals (N = 10), TD tubes present at Dudley Zoo during the sampling period but were never opened (N = 5), and several laboratory blanks with empty Tenax-filled TD tubes that remained in the lab and were analysed between sample measurements.

To collect *odour samples for bioassays*, we dabbed sterile gauze pads (viscose and polyester, Premier Healthcare & Hygiene Ltd, Great Britain) on the anogenital area for 20 sec. Samples were stored in 4 mL glass vials at −18 °C during sample collection and at −80 °C for storage. We used bioassay samples from the days included in the chemical dataset. Due to an unexpected 2.5-year delay in exporting samples from Zoo Dudley to Leipzig University, we collected additional samples for bioassays for comparison. They were collected over one complete cycle according to the original data set from two non-contracepted females aged 28 and 34 years living in one non-breeding group of five females and one sterilised male at the Wolfgang Köhler Primate Research Centre (WKPRC) in the Leipzig Zoo, Germany, in October and November 2025 following the same protocol.

### Hormone Analysis

Salivary progesterone concentrations were measured in the endocrinology laboratory of the German Primate Centre (Göttingen, Germany) in March 2025 for samples from Dudley and in December 2025 for samples from Leipzig using an enzyme immunoassay (EIA) based on the monoclonal antibody CL425 (Quidel clone no. 425, final purification by C. Munro, Davis, CA). Details of the analysis are described in Supplementary Material. Progesterone levels measured from saliva presumably reflect the hormonal state at the moment of sampling with only a slight time delay (20–30 min after appearance in blood (53)). Thus, we defined the periovulatory window as the two days immediately before the rise in progesterone level (two standard deviations above average of the five days before). Luteal phase samples were collected during the flat phase (after the progesterone rise) and follicular samples were collected during the increasing and maximal swelling stages (before the periovulatory window) (Fig. 1).

### GC–MS Analysis and Chemical Profiling

TD tube samples were shipped to the University of Leipzig immediately after collection and measured in September 2022 using a GCMS-TQ8040 (Gas Chromatograph GC-2010 Plus and a Triple Quadrupole Mass Spectrometer, Shimadzu, Kyoto, Japan) (see details in Supplementary Material). Chemical data were analysed using an established, semi-automated procedure (17) using AMDIS 2.73 (54) (see details in Supplementary Material). The final chemical data set included 200 substances for which we calculated relative intensities (peak area / sum of all peak areas x 100) to account for differences in sample intensities.

### Bioassays

Bioassays were conducted at Tacugama Chimpanzee Sanctuary (Freetown, Sierra Leone) from January to February 2026 (J.W.S.). Odour samples were presented to 18 male chimpanzees kept in different groups. Only 13 males (aged 8 to 42 years; mean ± SD: 18.5 ± 8.8) voluntarily participated in at least one test.

Thawed swab samples were placed in the setup (pre-cleaned with 70 % ethanol, wearing gloves) attached to the enclosure’s grid and allowing to present the samples in two open glass vials set 56 cm apart and facing the subjects. Samples remained in the setup for 5 min in presence of males. Male behaviour was video-recorded for subsequent analysis (see details of the setup and filming in Supplementary Material).

We conducted three types of tests in two-choice tasks in randomised order: samples from females in (1) follicular vs. periovulatory phase (N^males^= 16), (2) luteal vs. periovulatory phase (N^males^= 18) and (3) blank swab (sterile swab) vs. female sample (N^males^ = 19) (see more details in Supplementary Material). Males showed interest in samples only in 21 of 53 cases. As two males successfully participated twice under the same condition (luteal vs. periovulatory), their measurements were averaged to ensure independent observations for analysis. This resulted in the following sample sizes: four males for follicular vs. periovulatory tests, six males for luteal vs. periovulatory tests, and nine in blank tests. Due to low sample size, results are discussed descriptively.

Video analysis was conducted blindly to sample identity. We evaluated the number of investigation approaches and the total investigation duration. Investigating included (1) sniffing (bringing the nose within 4 cm of the sample or touching the mesh in front of the sample and sniffing at the finger within 3 sec); (2) licking (dipping the tongue or open mouth to the mesh in front of the sample). To control for an observer bias, a second observer coded twelve videos. An intra-class correlation (ICC) was calculated using the R package *irr* v. 0.84.1 (55), showing a high to excellent consistency between observers (ICC = 0.963 for number of investigations and ICC = 0.855 for investigation durations).

### Statistical Analysis

#### General statistical approach

As swelling stage might mediate the effect of menstrual cycle phase (Fig. 2), we needed to separate their direct and total (direct + mediated) effects. To do so, we conducted two analyses: (A) including both predictors in the same model to assess the direct effects for both; and (B) including only menstrual cycle phase to assess its total effect (see more details in Supplementary Material). All statistical analyses were conducted using R v. 4.4.2 (R Core Team, 2025).

**Fig. 2.**
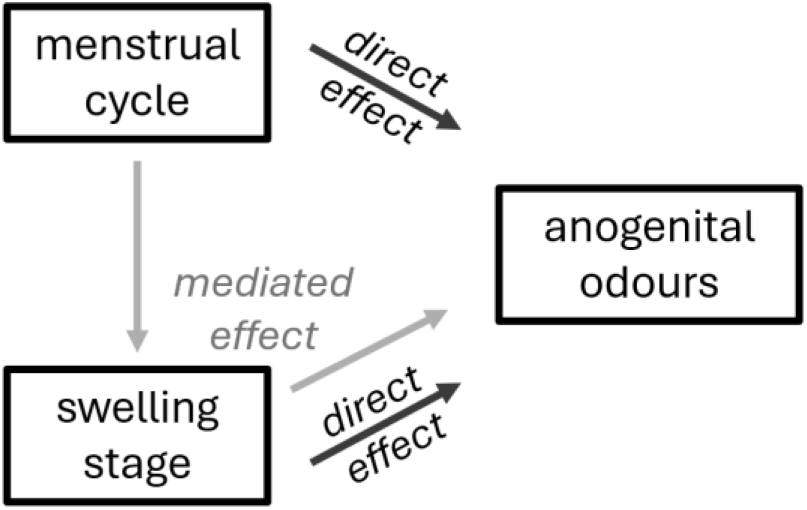
Simplified directed acyclic graph illustrating the direct effects of menstrual cycle phase and swelling stage on anogenital odours and the effect of menstrual cycle phase mediated by swelling stage.

#### Chemical composition

We fitted generalized linear mixed models (GLMMs) using the package *lme4* v. 2.0-1 (56) to investigate whether sexual swelling stage and menstrual cycle phase influence chemical composition of anogenital odours. For both variants (A) and (B), the vectorized data matrix was composed of all substances (N = 200) for all odour samples (N = 74), resulting in 14,800 substance intensities as response. Each model variant was fitted with two response transformations: (1) normalised and (2) log-transformed intensities. Normalisation centres all substance intensities to a mean of zero and a standard deviation of one, equalising the contribution of all substances and *increasing sensitivity on difference of substances among samples*. Log-transformation (log(x + 0.001) and arcsine) reduces the over-influence of the largest substances (57), but maintains the intensity differences of substances within samples and is *more sensitive to changes in condition-specific substances*.

Sexual swelling stage (four factor levels: ‘flat’, ‘increasing’, ‘maximal’, ‘decreasing’) and/or menstrual cycle phase (three factor levels: ‘follicular’, ‘periovulatory’, ‘luteal’), the enclosure type (three factor levels), and GC–MS batch (three factor levels) were fitted as fixed effects. Rows and columns (samples and substances) of the vectorized data matrix were fitted as random effects to account for pseudoreplication and heteroscedastic variance (58). Identity (ID, five factor levels) of the females of Zoo Dudley was used as additional random effect. We included all meaningful random interactions (swelling stages and cycle phases within ID) and fitted random interactions of sexual swelling stages and/or menstrual cycle phases within substances as test predictors and the random interaction of GC-MS batch within substances to get more reliable p-values. We estimated a random intercept for each substance × swelling stage/menstrual cycle phase combination, allowing to assess *differences between stages/phases for each substance*. Likelihood ratio tests (LRT (59)) were used to compare full models against null models lacking random interactions. Significant full-null model tests were followed by comparisons against reduced models excluding random interactions of each predictor separately, a method that has performed well for detecting effects of predictors on chemical substances in a simulation study (Weiß et al. in press).

We found no violations against the assumptions of normal distribution, homogeneity of residuals and residuals against fitted values for all models. Variance inflation factors, determined using the R package *car* v. 3.1-3 (60), did not exceed 2.0, indicating no collinearity issues.

#### Chemical profiles

We performed permutational analyses of variances (perMANOVA) in R package vegan v. 2.6-8 (61) to investigate the effects of menstrual cycle phase and swelling stage on whole chemical profiles of odours. As above, we used two variants (A) and (B) and two response transformations, normalised and log-transformed. The perMANOVA was performed on Bray–Curtis dissimilarities, with individual ID included as a strata term.

#### Substance identification

When a GLMM revealed significance, we investigated the differences for each substance between the levels of the test predictors. Following (62), substances with differences of 2.5 standard deviations above the average for all 200 substances were considered to be of particular importance. We tentatively identified these substances via library comparison using NIST14 Mass Spectral Library (National Institute of Standards and Technologies, Gaithersburg, MD, USA), accepting hits with a similarity index above 80. Suggested identifications were confirmed by comparing retention indices from NIST to those of a known alkane series measured on the same instrument using the same method.

## RESULTS

### Chemical Analyses

#### Chemical composition

When *both* variables were included (LRT full-null: χ2 = 65.9, df = 2, p < 0.001), we found a direct effect of swelling stage (LRT swelling: χ2 = 54.3, df = 1, p < 0.001), but not of menstrual cycle phase (LRT cycle: χ2 = 0, df = 1, p = 0.999) on chemical substances in the log-transformed model variant (Fig. 3). When the response was normalised, neither swelling stage nor menstrual cycle phase had a significant effect (LRT full-null: χ2 = 0, df = 2, p = 1).

**Fig. 3.**
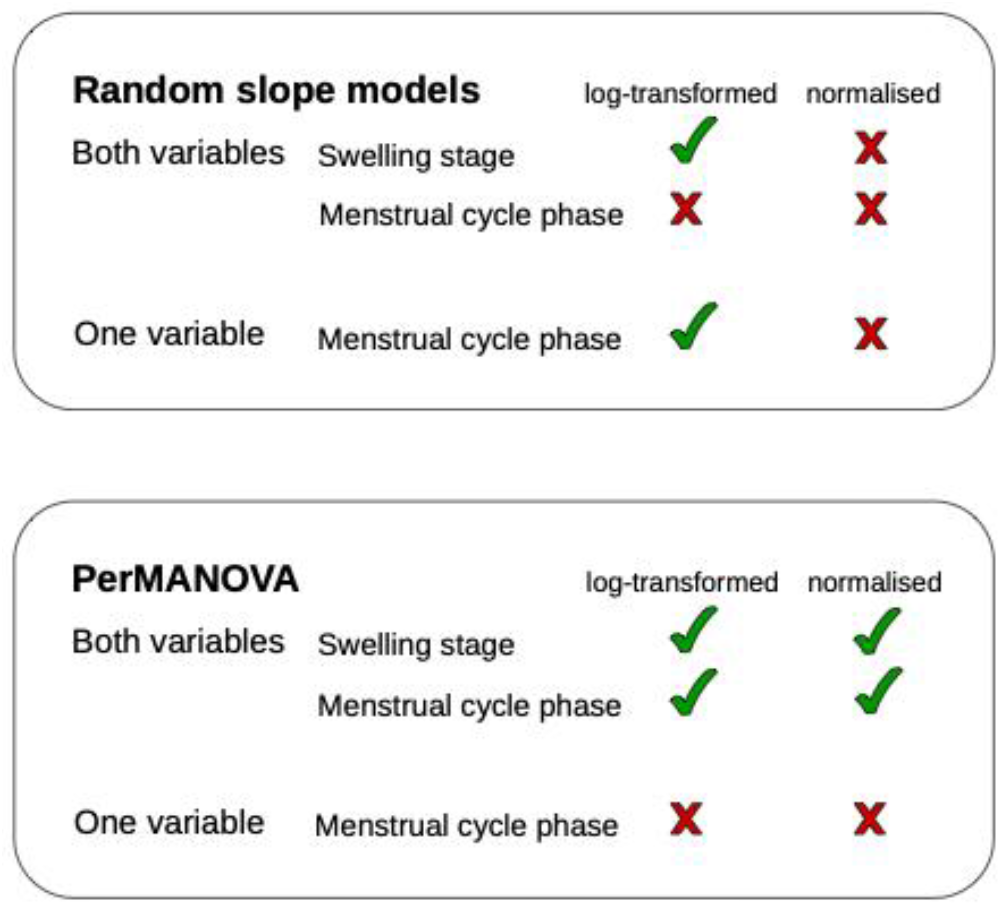
Summary of results. Green checks indicate a statistically significant effect; red crosses indicate the absence of significance.

When we included *only menstrual cycle phase*, it significantly affected chemical composition (LRT: χ2 = 11.6, df = 1, p < 0.001) with log-transformed, but not with normalised response (LRT: χ2 = 0, df = 1, p = 1). Based on the individual differences per substance with log-transformed response, five substances were found to be particularly affected by swelling stage and three substances by menstrual cycle phase (differences 2.5 SD > average; Supplementary Table S1, Fig. 4, 5).

**Fig. 4.**
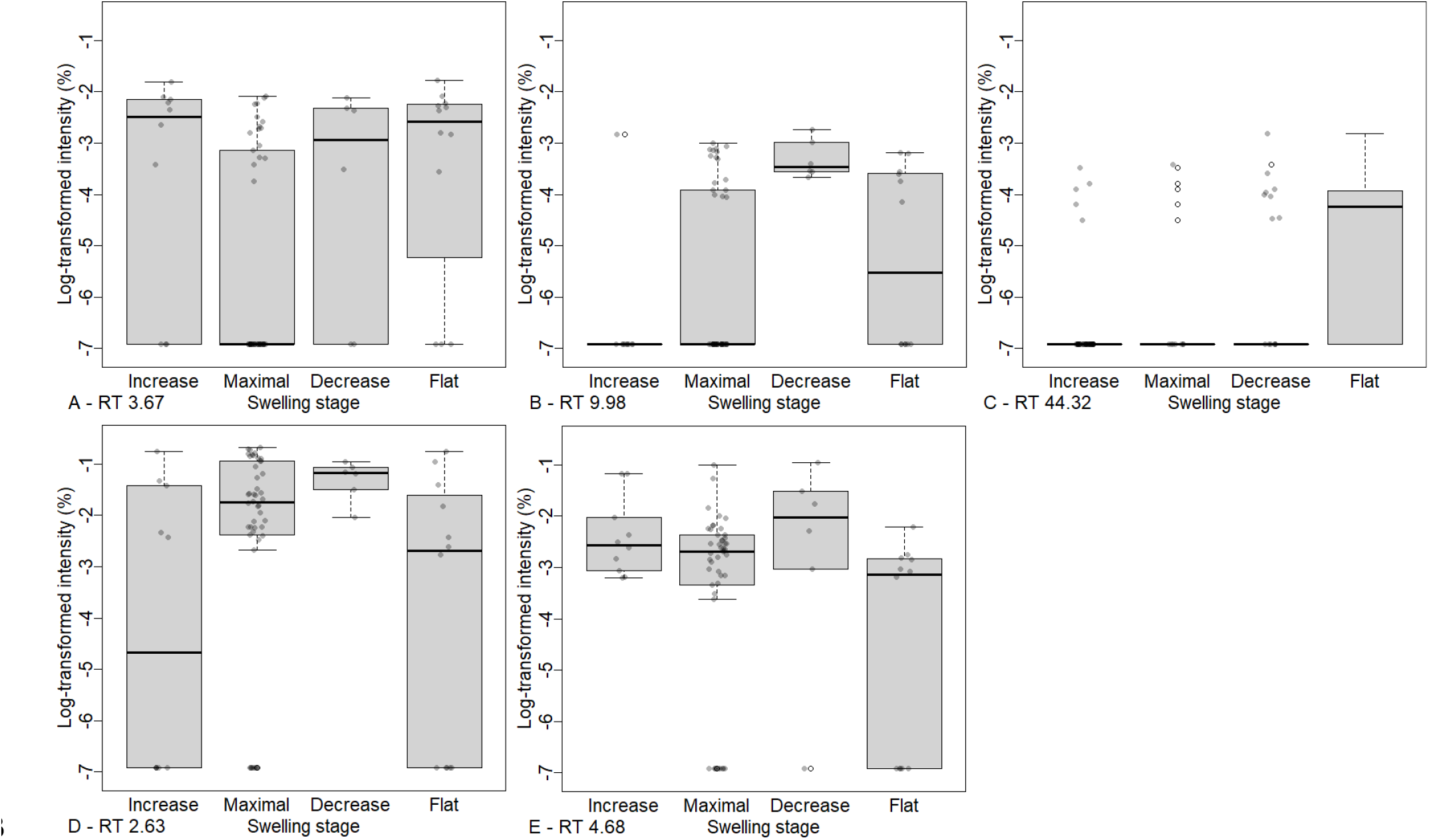
Substances most affected by swelling stage with log-transformed response, in descending order of effect strength. A: propanoic acid at retention time (RT) 3.67, B: undecane at RT 9.98, C: cholesterol at RT 44.32, D: acetic acid at RT 2.63, E: 2-methyl-heptane at RT 4.68. Boxplots show medians and first and third quartiles, grey dots represent raw data points.

**Fig. 5.**
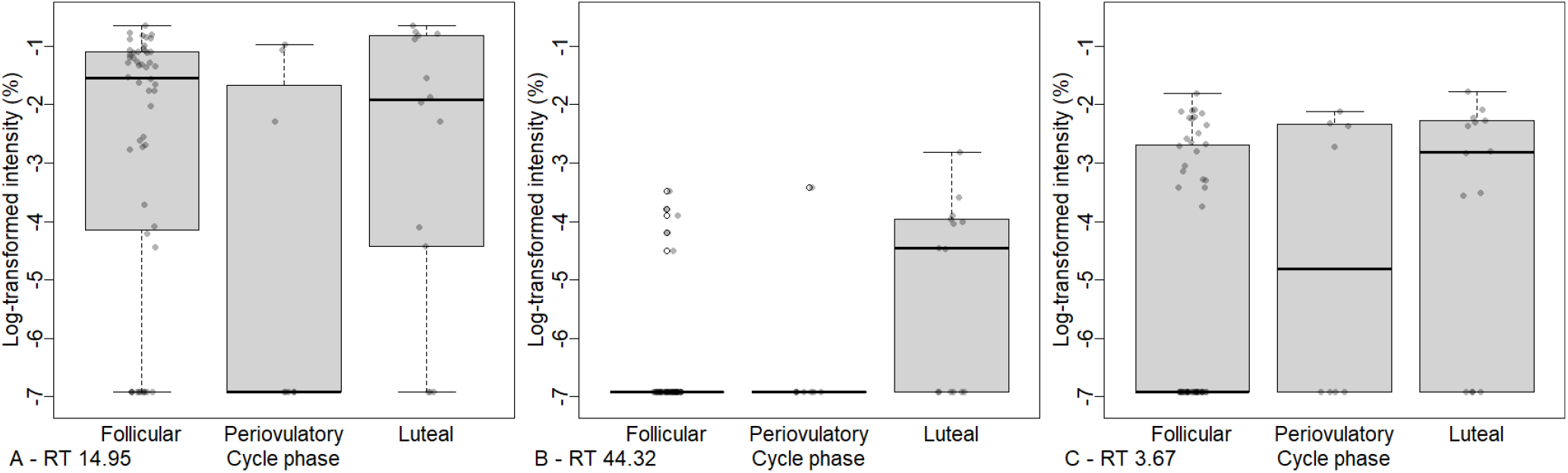
Substances most affected by menstrual cycle phase with log-transformed response, in descending order of effect strength. A: methenamine at RT 14.95, B: cholesterol at RT 44.32, C: propanoic acid at RT 3.67. Boxplots show medians and first and third quartiles, grey dots represent raw data points.

#### Chemical profiles

When *both* variables were included, swelling stage as well as menstrual cycle phase significantly affected whole chemical profiles with log-transformed (swelling: R^2^ = 0.09, F = 2.59, p = 0.004 and cycle: R^2^ = 0.05, F = 1.91, p = 0.046) and normalised response (swelling: R^2^ = 0.09, F = 2.20, p = 0.004 and cycle: R^2^ = 0.06, F = 2.14, p = 0.010). When *only menstrual cycle phase* was included, it affected the profiles neither with the log-transformed (R^2^ = 0.03, F = 1.21, p = 0.248) nor with the normalised response (R^2^ = 0.04, F = 1.34, p = 0.104).

### Bioassays

In 7 of 9 blank tests males investigated female samples more often than blanks, and in 6 of 9 blank tests investigation duration was longer for samples than for blanks. When presenting samples from follicular vs. periovulatory phase, only 1 of 4 males investigated the periovulatory sample more often and longer (N investigations: mean periovulatory ± SD: 1.3 ± 1.9 vs. mean follicular ± SD: 2.3 ± 2.6; investigation durations: mean periovulatory ± SD: 2.2 ± 3.9 sec vs. mean follicular ± SD: 3.5 ± 5.3 sec, Fig. 6). For the comparison of samples from luteal vs. periovulatory phase, 4 of 6 males investigated periovulatory samples more often (N investigations: mean periovulatory ± SD: 1.4 ± 1.4 vs. mean luteal ± SD: 1.1 ± 1.2) and 3 of 6 males investigated periovulatory samples longer (investigation durations: mean periovulatory ± SD: 1.7 ± 1.3 sec vs. mean luteal ± SD: 1.3 ± 1.3 sec).

**Fig. 6.**
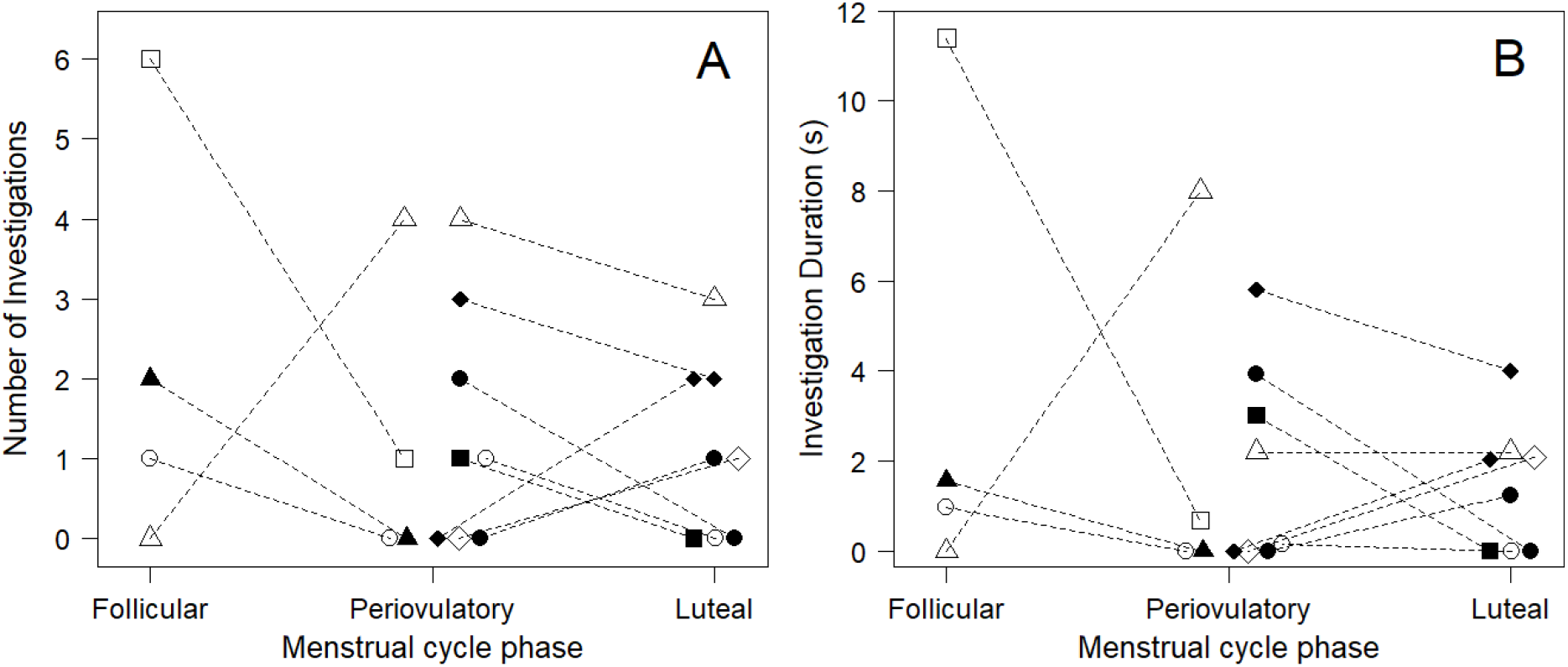
Investigation numbers (A) and durations (B) of odour samples from different menstrual cycle phases by males. Each symbol represents one male individual (two males are represented twice for luteal vs periovulatory, with a black circle and a black diamond), dotted lines connect data points of the same behavioural test run.

## DISCUSSION

Chemical composition of female anogenital odours varied across both sexual swelling stages and menstrual cycle phases. While the swelling stage had a significant direct effect on odours in most statistical analyses, an additional direct effect of the menstrual cycle phase extending beyond the effect of the swelling stage was subtle and could be revealed only in some analysis variants. Contrary to our expectations, we did not observe a male preference towards odours of fertile females. However, male interest toward the odour stimuli was generally too low to draw reliable conclusions about their ability to discriminate between odours from different phases of the menstrual cycle.

We hypothesized that olfactory cues provide more precise information about the timing of ovulation than the swelling size alone building on evidence that swelling size serves as a proxy for ovulation, combined with observations that male chimpanzees sniff at females during their maximal swelling and display increased mating interest during the periovulatory period (36,43,44). Our chemical results provide partial support for this hypothesis. First, we used models to examine whether individual substances change across swelling stages and/or menstrual cycle phases. Swelling stage had a direct significant effect on anogenital odours, corroborating our previous findings of odours from other body parts being affected by swelling stages (26). The menstrual cycle phase had a significant effect only when mediated by swelling stage but did not reveal a direct effect. The effects were significant only for log-transformed response, focused on differences between samples, but not for normalised response, focused on substance differences within samples.

Second, we used perMANOVA to examine whether whole chemical profiles change across swelling stages and/or menstrual cycle phases. Similar to the models, swelling stage had a direct effect on odour profiles, revealed in both log-transformed and normalised responses. Additionally, the menstrual cycle phase had a direct effect on odour profiles, extending beyond swelling and consistent across both response transformations. The overall chemical profiles, therefore, are influenced by both factors independently. These findings suggest that an olfactory signature of the menstrual cycle provides some *complementary* information about the timing of the periovulatory period. These olfactory changes, however, are minor (only 5-6 % of variance explained, marginally significant) compared to those related to the swelling stage (9 % of variance explained, highly significant).

The influence of swelling stage and menstrual cycle phase on odour was more consistent for the log-transformed response across all analyses. As the log-transformation allows substance intensities to differ in relation to the remaining chemical profile of a given sample, these findings suggest that fertility-related information could be encoded within odour chemical composition at a single time point. Consequently, males may be able to assess a female’s reproductive status through a single investigation, without a need for continuous odour monitoring. These findings are consistent with the dynamics of a fission-fusion society, where subgroup composition may change daily and males may lack consistent access to specific females. Furthermore, this would allow lower-ranking males, who may be socially precluded from consistent access to fertile females, to still obtain fertility cues during opportunistic encounters. However, in the analysis of the whole chemical profiles, the effects of swelling stage and menstrual cycle phase on odour were also found for the normalised response. Since with normalised response, affected substances might rather reflect changes over time, these findings indicate that additional fertility cues may be available to males continuously monitoring the same female over the course of her menstrual cycle.

In total, six chemical substances were affected by swelling stage and/or menstrual cycle phase (Fig. 4, 5). Five substances were affected by swelling stage, and two of them (excluding methenamine) were additionally affected by menstrual cycle phase, consistent with our finding that menstrual cycle phase does not have a direct effect on odours but is mediated by swelling stage.

Two of the affected substances, acetic acid and propionic acid, changed in levels mostly during the maximal swelling stage (acetic acid increased and propionic acid decreased), with propionic acid also increasing towards the periovulatory phase of the menstrual cycle. In previous studies, these aliphatic acids were found in vaginal secretions of chimpanzees (50,51), humans (63), and rhesus macaques (64,65), as well as in scent glands of common marmosets (66) (Supplementary Table S1). Furthermore, acetic and propionic acid, in a mixture together with other short-chain aliphatic acids, were reported to stimulate sexual behaviours in male rhesus macaques and were therefore referred to as sexual pheromones, or “copulins” (64,65), although these studies were later criticised (67). The amount of aliphatic acids in vaginal secretions, with acetic and propionic acids being the most prominent in all species, showed some variation across the cycle in humans and macaques, although the patterns were not consistent (63,65). In one chimpanzee study, a female with the highest levels of acetic and propionic acids was the only female followed by a male (51).

Cholesterol was more abundant during the flat swelling stage and in the luteal phase. Cholesterol is the precursor for steroid hormones such as oestradiol and progesterone (68), both of which are known to mediate swelling size. Specifically, increasing swelling is correlated with the rise of oestrogen levels in the follicular phase, whereas decreasing swelling follows a rise in progesterone, which inhibits the effects of oestrogen (33,48,49). As progesterone levels increase during the flat swelling stage and in the luteal phase (33,68), the increase in cholesterol levels observed during this period may reflect the need for progesterone synthesis. In line with this, a previous study in chimpanzees demonstrated cyclic variation in cholesterol-derived substances, which were more abundant during the increasing and decreasing swelling stages compared to the maximal swelling and flat stages (26), probably reflecting cyclic patterns of processing and utilization of cholesterol for biosynthesis of steroid hormones.

Two other affected substances were alkanes (2-methyl-heptane and undecane) that had previously been found in human breath, saliva, skin, or other tissues (69,70) and, in the case of undecane, in the sternal scent gland of mandrills (71,72) (Supplementary Table S1). While they had not been reported in a fertility-related context, their changes across the menstrual cycle may reflect broad physiological changes associated with cyclic hormonal changes (73). Finally, methenamine, showing the lowest level during the periovulatory phase, has non-mammalian origins and is used as a medication, as well as in production of plastics and cosmetics. Methenamine was previously found in anogenital odours of common marmosets, where its levels were higher in nulliparous females compared to females who gave birth to infants already (17) and was discussed as potential contamination released through the genital secretions.

Subtle changes in chemical profiles of female odours across menstrual cycle phases, extending beyond swelling stages, could potentially provide additional information on fertility to individual males. While visual signals of swelling size are broadcast to all males, dominant males are likely able to approach and smell the female anogenital region more often. This is corroborated by an earlier finding where the dominant male, when having the choice among several maximally swollen females, preferred the fertile female rather than that with the biggest swelling (36). These findings align with the graded signal hypothesis that proposes that swellings are probabilistic signals of the timing of ovulation (74). According to this hypothesis, information on ovulation probability allows dominant males to monopolize a female during peak fertility, while still allowing other males to achieve copulations during periods of lower ovulation probability. At the same time, this enables females to balance the benefits of confusing paternity (to reduce the risk of infanticide) with biasing paternity towards the dominant male (to increase paternal care and the probability of obtaining ‘good genes’ for their offspring) (74,75). In line with this argumentation, an earlier study showed that, despite clear male dominance, female chimpanzees effectively exhibit mate choice by being more proceptive and less resistant to preferred males during the fertile period, resulting in these males having higher mating success than other males (38).

Similar findings in support of the graded-signal hypothesis and multimodal fertility communication were demonstrated for olive baboons, a species with a social system similar to that of chimpanzees, where females live in multi male-multi female groups and mate with multiple males. In this species, too, sexual swellings were shown to be complemented by more precise information on the timing of fertility window provided by vaginal odours (25). Similar to chimpanzees, sniffing and copulation rates by dominant male baboons increased during fertile days, suggesting the role of olfactory cues accessible to males at closer range (28,76). There is also some tentative evidence of urine odour differences between fertile and non-fertile phases in Japanese macaques, another species with a multi male-multi female social system, multiple female mating, and visual signals of fertility (24,77).

Olfactory fertility cues, however, are not necessarily accompanied by visual signals, as is evident from research on common marmosets, who lack visual fertility signals but nevertheless exhibit clear fertility-related odour changes (17). Another species lacking obvious visual fertility signals are modern humans. Chimpanzees are closely related to humans phylogenetically and share many aspects of social organization and menstrual-cycle physiology with them. It is plausible that modern women may share olfactory fertility cues with chimpanzees, too. However, research on humans shows mixed evidence: while some studies have reported evidence for olfactory fertility cues (78–80), others failed to replicate these findings (81–84). It thus remains unclear whether olfactory cues of fertility have been lost, together with visual fertility cues, over the course of human evolution. Thus, the existence of olfactory fertility cues might not depend on the presence or absence of visual signals, but rather on species-specific costs and benefits of communicating fertility via specific sensory modalities.

It remains unclear whether odour changes detected by our chemical analyses are perceivable and relevant to males, because the males participating in bioassays exhibited insufficient interest in the odour samples to allow for a meaningful analysis of preference. In addition to low interest in general, successful trials did not reveal any systematic preferences for odours from fertile females. It is possible that olfactory information is only relevant as part of multimodal cues (combined with visual signals and/or behavioural cues). However, it is more likely that the lack of significant response is related to methodological limitations that resulted in low motivation and lack of focus in male subjects. The males were often distracted by external factors, such as vocalizations and aggressive interactions of animals in the neighbouring enclosures and the calls of wild chimpanzees in the surrounding forest. In some cases, males were fed directly before testing, which caused inactivity, or found leftover food in the enclosures, which distracted them from the experimental setting. The experimental setup was novel to the animals, and some of them focused on the setup itself instead of odour samples, chewing or tearing at the setup. It is also possible that the lack of previous positive experience of participating in bioassays lowered their motivation to investigate the samples. Consistent with this, the rate of no-sniffing was significantly higher at the sanctuary, where 32 of 53 trials resulted in a failure to sniff (60 %), compared to the Leipzig Zoo pilot study, where chimpanzees are frequently rewarded for test participation and only 5 out of 30 trials resulted in a failure to sniff (17 %). Finally, the volatile composition of the samples may have been compromised by inconsistent storage during fieldwork, including an eight-hour period where temperatures reached 1.5 °C. However, this likely had a negligible impact, as freezing has been shown not to affect odour perception (85) and the interest in samples did not differ between samples from Leipzig and samples from Dudley.

## Conclusions

We predicted that (1) the chemical composition of female odours will vary across menstrual cycle phases in addition to the variation between sexual swelling stages and provide more precise information on fertility than the swelling alone, and that (2) males will be able to detect these differences. Our chemical analysis partially supported the first prediction. In addition to the pronounced olfactory signature of sexual swelling, changes in whole chemical profiles associated with menstrual cycle phases provided some complementary information about the timing of the periovulatory period. These olfactory changes, however, are minor compared to those related to swelling stages. The second prediction could not be confirmed as there was no indication of preferences towards odours from the periovulatory period. Additionally, low male interest toward the odour stimuli does not allow to draw any conclusion about their ability to discriminate between odours from different phases of the menstrual cycle.

Overall, our findings support the existence of a multimodal fertility cue in chimpanzees. The visual signal of swelling size is mirrored by olfactory changes in anogenital odours. In addition, subtle changes in overall chemical profiles also showed direct effects of menstrual cycle phase complementing them and seem to augment previous findings of slight increases in swelling size that indicate approaching ovulation (36). Direct effects of menstrual cycle phase, however, appear to be modest compared to those associated with swelling stages.

## Supporting information

Supplementary Material - Original Data

## ACKNOWLEDGEMENTS

We thank Jodie Dryden, Stacey Evans, Harley Hunt, Juliet Williamson, Emma Gardiner and Laura Pickersgill for their support in collecting odour samples at Zoo Dudley. We are especially grateful to Stefanie Bley, Jo Frommen, and Jerome Micheletta for the extraordinary effort involved in organizing sample transport to Germany.

We are grateful to the team at the WKPRC for allowing us to test the bioassay setup and collect new reference samples. In particular, we thank Daniel Haun and Daniel Hanus for granting permission, all animal caretakers of Leipzig Zoo for their valuable support, especially Daniel Geissler, René Berger and Hannah Petschauer for the primary coordination, and Raik Pieszek for assistance with the design of the setup.

We thank all supporters involved in the bioassays at Tacugama. We are grateful to Bala Asakamaram and Nicolas Quiroz for permission, to Sophie Collier, Philip Guy, Pastor Kamara, Camila Camargo, Timone Schoonhoven, and Asami Kabasawa for organisational and logistical support, to all caretakers for their assistance — especially Daniel, Albert, Yomin, Wilson, Abu, Saido und Mama P — and to Josephine Kalbitz for support with the study planning. We thank Antonia Krüger and the MS Core Facility at Leipzig University for support with measurements and chemical profiling.

Finally, we are grateful to Verena Behringer and the endocrinology lab of the German Primate Center for conducting the hormone analyses. Equipment for sample transport and refrigerated storage was generously provided by delta T GmbH. We sincerely thank Andreas Schön for the excellent advice and support.

## STATEMENTS

### Ethics

The study was ethically approved by the European Association of Zoos and Aquaria (EAZA), especially the EAZA Ex situ Programmes (EEPs) for Chimpanzees as well as sample collection by the Zoo Dudley, UK. Samples were obtained and transported under CITES export and import regulations in compliance with relevant national and international regulations (export permit no: 24GBEXPR4E6RF; import permit no: DE-E-05276/24) and were transferred under import permit issued by the Saxon State Ministry for Social Affairs and Social Cohesion.

Sample collection at the Wolfgang Köhler Primate Research Centre (WKPRC) in the Leipzig Zoo, Germany was in accordance with the legal requirements of Germany and approved by the Ethics committee of the Leipzig Zoo and the Department of Comparative Cultural Psychology, Max Planck Institute for Evolutionary Anthropology in Leipzig, Germany.

Protocols for bioassays were reviewed and ethically approved by the veterinary department of the Tacugama Chimpanzee Sanctuary, Sierra Leone.

### Data, code and materials

The datasets supporting this article have been uploaded as part of the supplementary material.

