## Supplementary Material - Additional Information for "Multimodal fertility cues in chimpanzees: How body odours complement sexual swellings"

### METHODS

#### Hormone Analysis

Progesterone–horseradish peroxidase was used as a conjugate (1). CL425 was raised against 4-pregnen-11-ol-3, 20-dione hemisuccinate:bovine serum albumin and shows broad cross-reactivity with progesterone (100 %) and its metabolites (2). The assay successfully applied to capture cyclical variation serum samples across a range of mammalian species (3), including primates (1), supporting its suitability for assessing reproductive state. Analytical validation in serum has demonstrated high sensitivity (90 % binding = 50 pg mL−1), parallelism between standard curves and serial dilutions of samples, and high precision (intra- and inter-assay coefficients of variation < 10 % and < 15 %, respectively) (4). Parallelism between serially diluted saliva samples and the standard curve was confirmed.

For the assay, saliva samples were diluted with an assay buffer at dilutions of undiluted, 1:2 or 1:3 depending on hormone concentration. Plates were washed prior to use, and 50 µL of either diluted sample or progesterone standard was added to each well, followed by 50 µL of horseradish peroxidase conjugated and 50 µL of antibody solution. After overnight incubation at 4 °C, plates were washed and 100 µL of tetramethylbenzidine (TMB) substrate was added. The reaction proceeded for 45 minutes in the dark at room temperature on a plate shaker. The enzymatic reaction was stopped by adding 50 µL of 2 M sulfuric acid, and optical density was measured at 450 nm with a 630 nm reference filter using a microplate reader. Results are expressed in pg/mL.

All samples and standards were measured in duplicate. Samples with a coefficient of variation exceeding 10 % or falling outside the linear range of the assay were reanalysed. Quality control (QC) samples with known high and low progesterone concentrations were included in each assay run. Inter-assay coefficients of variation (N = 8) were 10.1 % (high QC) and 5.4 % (low QC).

#### GC–MS Analysis and Chemical Profiling

Using a thermal desorption system TD-20 (Shimadzu, Kyoto, Japan), samples were desorbed at 250 °C for 8 min to a Tenax TA filled cold trap (−20 °C) with a helium flow of 60 mL/min in split mode (split ratio 5). The injection into the gas chromatograph was performed at a pressure of 140.2 kPa by heating the cold trap to 250 °C for 1 min. One-dimensional chromatography was done using two consecutively connected columns (Rxi-1ms: 30 m length, 0.25 mm ID, 0.25 µm film, and SGE Analytical Science BPX50: 2 m length, 0.15 mm ID, 0.15 µm film, Restek GmbH, Bad Homburg vor der Höhe, Germany) with helium 5.0 as carrier gas at a flow rate of 1.58 mL/min. The temperature program started at 35 °C for 0.5 min and ended at 320 °C for 25 min after a rise of 6 °C/min. Electron impact ionization was conducted at 70 eV and a source temperature of 220 °C. Mass spectra were acquired over a scan range of m/z 30 – 500.

Chemical data were analysed using an established, semi-automated procedure (5) using AMDIS 2.73 (6) for peak picking to create an aligned list of substances present in the samples and integrating substances using GCMSsolution v. 4.20 (Shimadzu, Kyoto, Japan) to get substance intensities. The resulting data included only substances with a signal-to-noise ratio (S/N) above 1.0 and were cleaned by removing incorrectly integrated substances outside the desired retention time range, that occurred in less than 5 % of the samples and that were more intense in blank samples than animal samples or were known contaminations.

**Bioassays**

We developed and tested the bioassay setup at the WKPRC with male chimpanzees trained to these sets of tests. However, the WKPRC at that time had only 2 adult males available. Therefore, the preference tests using swab samples were conducted at Tacugama Chimpanzee Sanctuary (Freetown, Sierra Leone).

We transported frozen swab samples to Tacugama in a vacuum bag with cooling packs and phase-change elements (delta T GmbH, Fernwald, Germany) and stored them at −6.18 °C (± SD: 2.8 °C). A temperature tracker (ThermoScan 2S, delta T) continuously recorded the temperatures and detected one rise above the freezing point to 1.5 °C for approx. 8 h during a power cut which we considered not critical for the testing success, since repeated thawing-cycles do not seem to have a major impact on sample quality (7). Before testing, we thawed the samples (mean ± SD: 2 h 24 min ± 42 min).

The self-built setup made from acrylic glass (Fig. S1) was secured to the enclosure's grids, with additional wire mesh fitted in front of it. Opened glass vials containing the samples were inserted into two tubes accessible from the outside, so that the sample opening was positioned behind a mesh directly at the front opening facing the chimpanzee. The samples were spaced 56 cm apart from each other.

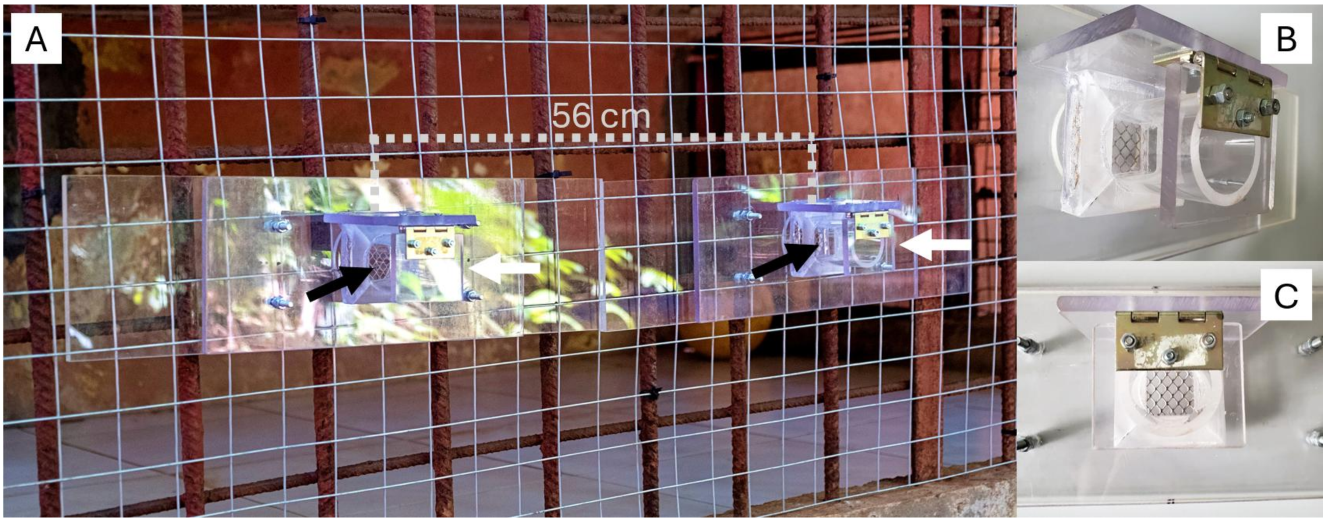


**Fig. S1.** Setup for bioassays. A: Original positioning during a trial, with two tubes (white arrows, 45 mm diameter, 56 cm apart from each other) for holding the opened glass vials with odour samples directly behind a mesh (black arrows). B: Lateral close-up view and C: frontal close-up view.

Samples were handled wearing gloves at all times and were left in the setup for five minutes from the moment the males were able to approach it. The behavioural tests were video recorded and afterwards analysed using the video recordings. For filming, we used at least one camera positioned directly in front of the setup, showing both samples simultaneously. In addition, depending on the enclosure characteristics, we filmed a side view from a diagonal front or diagonal rear angle. The person handling the samples and cameras (J.W.S.) was blind to the sample identity and kept out of sight and/or concealed any facial expressions by wearing sunglasses and a baseball cap during the tests.

The two samples within a single preference test always originated from different females to avoid the unnatural situation of encountering odours from the same female from two different time points. None of the males was presented with the same combination of female samples twice and at least four days of rest period were between tests involving the same male. Within each preference test, two samples from Dudley or two samples from Leipzig were presented, but never a mixture of the two locations. Due to the limited number of samples, odour samples were repeatedly used, but samples with the same number of uses were always paired together and the rate of investigations did not change between first usage and repeated usage trials. Usually, males were separated in an enclosure for the tests. For three groups a separation of single males was not possible and the setup was presented to a group of individuals. Males did not receive a reward for participating in the bioassays, so as not to induce learned behaviour.

### Statistical Analysis

*Additional explanation of statistical approach*

If both predictors are included together (A), the estimated effect for each of them is investigated while controlling for the other predictor, i.e. we unravel the direct effect of one variable in addition to what the other predictor already explains. If only one predictor is included at a time (B), the effect of this predictor is estimated including the variance of the other variable, because it is not controlled for in the analysis. See (8,9) on mediating effects in causal inference.

**Table S1.** Substances most affected by swelling stages and/or menstrual cycle phases with tentative substance identification (ID), average retention time (RT), similarity index (SI) indicating how well measured mass spectrum and reference spectrum matched, corresponding CAS number, molecular weight (MW) and substance class category. A numerical ranking of substances affected by swelling stages (Top Swell) and menstrual cycle phases (Top Cyc) represent the strength of the effects (1 = strongest effect) and the differences between swelling stages and menstrual cycle phases are given in brackets (threshold of average + 2.5 SD: 0.58 for swelling stages and 0.27 for menstrual cycle phases). Some example references are given for studies on primates including humans that found the same substance.

| Substance ID | RT | SI | CAS | MW | Substance class | Top Swell | Top Cyc | References |
|---|---|---|---|---|---|---|---|---|
| Acetic acid | 2.63 | 97 | 64-19-7 | 60 | carboxylic acid | 4 (0.66) | | - Vaginal secretions of chimpanzees (10,11), humans (12), and rhesus macaques (13,14)<br>- Circumgenital gland secretions of female common marmosets (15)<br>- Sternal gland of mandrills (*Mandrillus sphinx*) (16)<br>- Faeces, urine, breath, skin, milk and saliva of humans (reviewed in (17)) |
| Propanoic acid | 3.67 | 92 | 79-09-4 | 74 | carboxylic acid | 1 (0.9) | 3 (0.32) | - Vaginal secretions of chimpanzees (10,11), humans (12), and rhesus macaques (13,14) |

| | | | | | | | | |
|---|---|---|---|---|---|---|---|---|
| | | | | | | | | - Circumgenital gland of female common marmosets (15)<br>- Faeces, urine, breath, skin, milk and saliva of humans (reviewed in (17))21/05/2026 09:14:00 |
| Heptane, 2-methyl- | 4.68 | 96 | 592-27-8 | 114 | alkane | 5 (0.63) | | - Breath and saliva of humans (reviewed in (17)) |
| Undecane | 9.98 | 92 | 1120-21-4 | 156 | alkane hydrocarbon | 2 (0.66) | | - Sternal scent gland of mandrills (16,18)<br>- Human axillary sweat (19), faeces, breath, skin, milk and saliva (reviewed in (17)) |
| Methenamine | 14.95 | 97 | 100-97-0 | 140 | | | 1 (0.36) | - Breath of humans (reviewed in (17))<br>- Anogenital skin of common marmosets (5) |
| Cholesterol | 44.32 | 84 | 57-88-5 | 386 | sterol | 3 (0.66) | 2 (0.35) | - Glandular secretions of ringtailed lemurs (20) |
